# Automated image processing to support the analysis of between-year transmission of *Leptosphaeria maculans* in field conditions

**DOI:** 10.1101/488890

**Authors:** Lydia Bousset, Marcellino Palerme, Melen Leclerc, Nicolas Parisey

## Abstract

Understanding the transmission of inoculum between periods where the host plants are present is central for predicting the development of plant diseases and optimising mitigation strategies. However, the production at the end of the growing period, the survival during the intercrop period, and the emergence or emission of inoculum after sowing or planting can be highly variable, difficult to assess and generally inferred indirectly from symptoms data. As a result, there is a lack of large data sets which is a major brake for the study of these epidemiological processes. Here we focus on *Leptosphaeria maculans* that causes the black leg of oilseed rape. After having infected leaves, at early stages of the plant, and migrating into the stem, it causes a basal stem canker before harvest. It then survives on stubble left in the field from which ascospores are emitted at the beginning of the next growing period. In this study we first developed an image processing framework to estimate the density of fruiting bodies produced on stubble. Then, we used this framework to analyse automatically a large number of stems collected in oilseed rape fields among a cultivated area. Having performed a quality assessment of the processing chain we used the output data to investigate how the potential level of inoculum may change with the source field, the considered year and the stem canker severity at harvest. Besides the insights gain into the blackleg of oilseed rape, this work shows how image-based phenotyping may support epidemiological studies by increasing substantially the precision of high throughput disease data.

## 1 Introduction

In agro-ecosystems many plant diseases have cyclic epidemics and their dynamics are highly influenced by both temporal and spatial discontinuities either induced by the climate (e.g. seasonality) or by human actions (e.g. sowing and harvesting) (Bousset and Chèvre, 2013; Hamelin et al., 2011). From an epidemiological point of view, the space-time partitioning of host-crops, and the space-time pathogen transmission among the cultivated area and between growing seasons are main determinants of both invasion and persistence of plant diseases (Bousset and Chèvre, 2013; Gilligan, 2002; Hamelin et al., 2011). Within these determinants, the between-year (or growing period) transmission of the pathogen, that gives the level of primary inoculum at the beginning of the next growing season, remains particularly difficult to predict and estimate (Bailey et al., 2004; Bousset et al., 2015). However, this temporal dispersal of the pathogen is an important step to design and predict the level of success of mitigation strategies, either based on the use of resistant varieties (Marcroft et al., 2004b), biocontrol agents (Bailey et al., 2004) or preventive management of crop residues (Wherrett et al., 2003).

In that context, the necessary epidemic data are generally difficult, costly and time-consuming to acquire (Bousset et al., 2015, 2016). Furthermore, as the direct quantification of the pathogen (or inoculum) and the infectiousness status of hosts plants (Susceptible, Latent, Infectious, Removed) are still challenging for plant diseases, they are indirectly inferred from pathology status data (Leclerc et al., 2014), often insufficient for testing mechanistic models without resorting to time-consuming Bayesian methods (Bousset et al., 2015). In this context, developing high-throughput and high-precision phenotyping methods appears as a main challenge for plant disease pathology and epidemiology (Bousset et al., 2016; Simko et al., 2016). For instance, as already shown by several authors, image-based phenotyping can be useful for detecting and quantifying disease symptoms (Camargo and Smith, 2009; Mahlein, 2016) and automated image processing enables one to expand substantially the throughput of disease data (Karisto et al., 2018; Stewart et al., 2016) which may feed modelling approaches and support empirical studies.

Depending on the pathogen, the inoculum that can survive between the periods where the host-crop is present can be propagules (e.g. sclerotia, oospores) lying into the soil or pathogen structures (e.g. mycelia) surviving on host (or alternative hosts) debris within or above the soil. In the particular case of stubble-borne diseases, after some delay during which fruiting bodies are formed and when suitable environmental conditions occur, infectious spores are released and dispersed around the stubble inoculum-source. Then, besides trapping the suitable aerial spores, one can use the number of fruiting bodies as a proxy of the level of the source of inoculum (Bousset et al., 2015). One reason that has slow down the researches in this direction is the technical difficulty of estimating fruiting bodies numbers on a large number of emitting debris. Direct enumeration can be achieved for large sclerotia of *Sclerotinia sclerotiorum* (Taylor et al., 2018) but becomes impractical for microsclerotes of *Ramolispora sorghi* (Brady et al., 2011). Similarly some authors have been able to estimate perithecia of *Mycosporella fijiensis* on leaves (Burt et al., 1999) or pseudothecia of *Leptosphaeria maculans* on stubble (Lô-Pelzer et al., 2009b), but, without the use of automated phenotyping methods the observation time, and the tediousness, limit the number of samples that can be processed.

The simplest use of digital images for counting fruiting bodies is to provide a way to postpone the assessments, thus enabling to process more samples than would be possible in real time. On pictures of excised leaf discs infected with *R. sorghi*, Brady et al. (2011) manually marked the microsclerotia before counting them and retrieving their sizes from numbers of pixels. This could be regarded as a slight improvement to excising the necrotic area and measuring it by tracing the shape of the cut section on to graph paper and later counting the numbers of perithecia of *M. fijiensis* under the microscope (Burt et al., 1999). A more advanced use of imaging is to use common processing methods and algorithms, generally implemented in open software such as ImageJ (Schindelin et al., 2015) or Ilastik (Sommer et al., 2011), to segment regions or objects. Nowadays, there are numerous instances of such use of imaging in phytopathology, e.g. detection and counting of microsclerotes of *Calonectria pseudonaviculata* (Yang and Hong, 2018) or quantification of lesion size and number of *Zymoseptoria tritici* on wheat leaves (Karisto et al., 2018; Stewart et al., 2016). At this stage, it is important to evaluate the quality of the segmentation method (supervised or unsupervised), for instance by assessing the discrepancy between the processed images and a set of images that were annotated by an expert (i.e. ground truth). When the image processing framework is reliable enough one could use it as a pipeline for generating a large number of automatic measurements.

In this study we concentrate on *L. maculans* that causes blackleg on oilseed rape crops and has a substantial economic impact worldwide. Epidemics are initiated early in the growing season by stubble-borne spores that can spread between fields among the landscape and the estimation of the production of inoculum sources, or the level of inoculum that is transmitted between growing seasons, is challenging and requires dense data (Bousset et al., 2015). We begin by presenting how we collected oilseed rape stubble left after harvest in a cultivation area during six years in order to assess stubble-borne production on inoculum between growing seasons. Then, we develop an image-prosessing framework to segment fruiting bodies produced on collected stubble. After having assessed the quality of the segmentation method we used it to process automatically a large number of images, and thus collected stubble. It allowed us to produce many pathological data which were used to evaluate how fruiting bodies’ production changes between years and fields and how the level of production can be linked with the blackleg severity after harvest. We finish by discussing the interest of our work for plant disease epidemiology and presenting some perspectives. An overview of our proposed approach is provided (Fig. 1, see also Fig S2).

**Figure 1:**
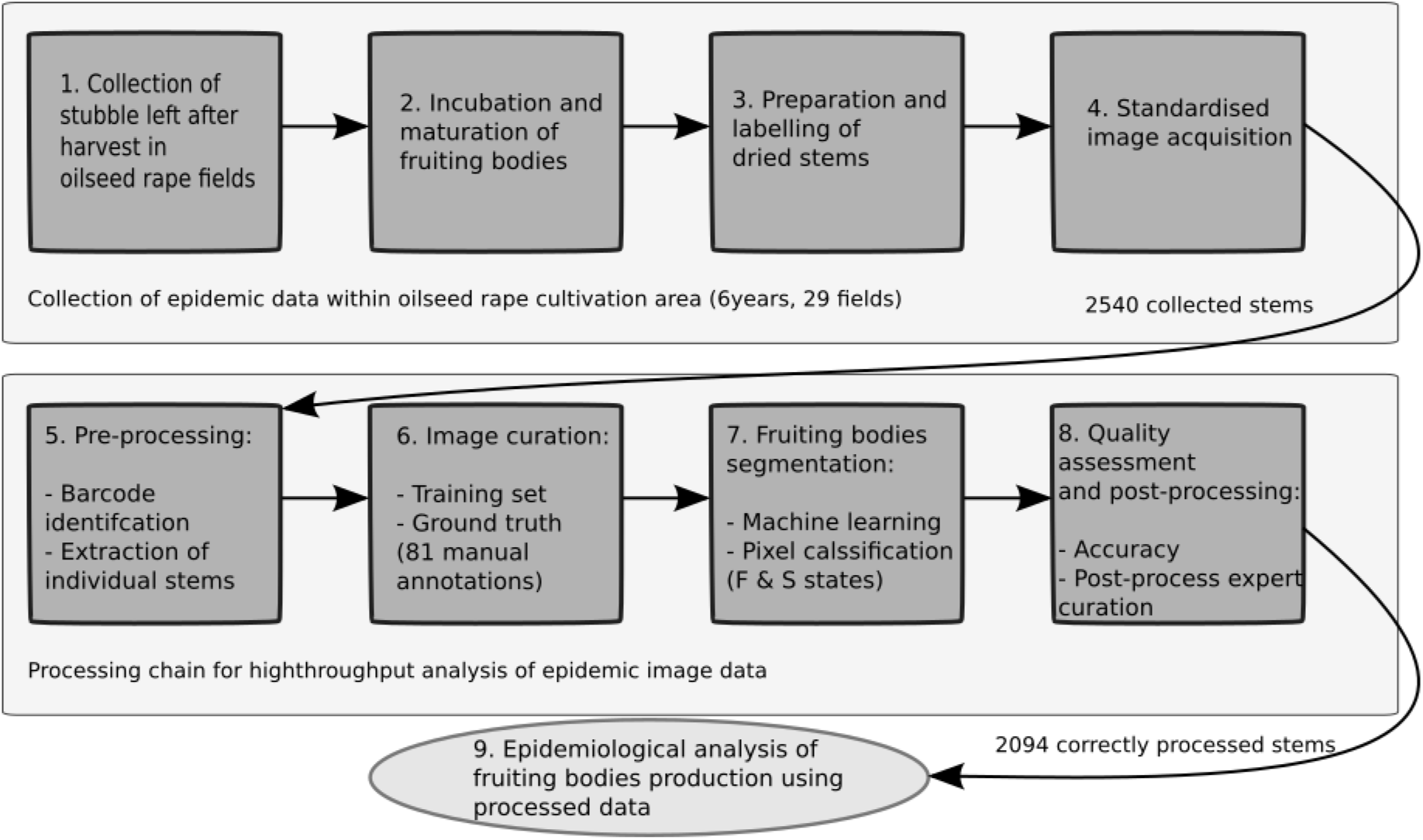
Successive steps in the automated image processing pipelines to support high-throughput epidemiological analyses.

## 2 Materials and Methods

### 2.1 Pathosystem

*Leptosphaeria maculans* causes stem canker on Brassica species (West et al., 2001). Epidemics are initiated in autumn and leaf spots are observed from autumn to early spring. Stem cankers develop from spring to summer, up to the time of harvest, following systemic growth of fungal hyphae from leaf spots to the leaf petiole through xylem vessels, and subsequently to the stem base. The fungus can survive as hyphae in crop stubble, forming two kinds of fruiting bodies: pycnidia and pseudothecia. Pseudothecia can only be formed by sexual reproduction if isolates of opposite mating types co-occur in the same oilseed rape stem. Spores produced in pseudothecia are, respectively, conidia (pycnidiospores) passively rain splashed short distances and ascospores actively ejected, and wind dispersed (Bousset et al., 2015; Marcroft et al., 2004a; Savage et al., 2013). Infected stubble ensures the carry-over of the fungus from one season to the next (Bousset et al., 2018; Lô-Pelzer et al., 2009b; Marcroft et al., 2004b,a), and serves as the main source of inoculum. The severity of blackleg in the plant at maturity was related to the inoculum later produced from the stubble (Lô-Pelzer et al., 2009b; McGee and Emmett, 1977). The formation of pseudothecia and hence the discharge of primary inoculum was influenced by the genotype of the infected stubble (Marcroft et al., 2004b) and by chemical treatment (Wherrett et al., 2003). The quantification of inoculum was achieved either by direct counting under the microscope (Lô-Pelzer et al., 2009b) or by visually recording pseudothecial density in classes from 0 to 100% (by 10% increment) of the stubble surface area infested with pseudothecia (Marcroft et al., 2004b; Wherrett et al., 2003). As the area occupied by pseudothecia was correlated with pseudothecia numbers (Lô-Pelzer et al., 2009b), automated detection on pictures could be used for the quantification

### 2.2 Field data collection and corresponding climate covariates

Oilseed rape stems were collected in a total of 29 farmers oilseed rape fields located near Le Rheu (48.1° N, 1.5° W), in Brittany, France, from June 2013 to June 2016, 2-3 weeks before harvest (Table 1) (Bousset et al., 2015). Stem canker severity was assessed on a 1 to 6 scale (Delourme et al., 2014) as follows: S1 = no disease, S2 = 1-25%, S3 = 26-50%, S4 = 51-75%, S5 = 76-99%, S6= 100% of crown cross section cankered. Following assessment of canker severity on 9901 stems, 2540 pieces of stubble comprising the crown and upper 10 cm were selected. From each of the fields, a maximum of 30 stems in each of the 6 severity classes was kept. This sampling was constrained by the availability of stems, as the distribution of stems into canker severity classes depends on the overall disease severity in the field of origin (Lô-Pelzer et al., 2009a). To keep track of stem canker severity of stem pieces throughout the experiment, two 5 mm diameter holes were drilled, and stem pieces were grouped by 5 on two wooden BBQ sticks painted in blue and labelled with a barcode (Fig. S1AB).

**Table 1:**
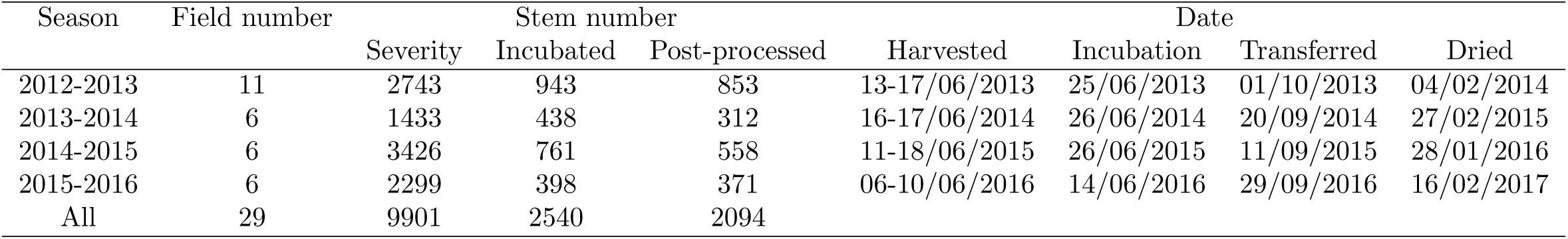
Numbers of stems assessed for canker severity, selected for incubation, kept after post-processing. Timing of stem harvest, start of incubation, transfer on fiend plots and end of incubation by drying.

Over summer, each selected stubble was matured outside at INRA Le Rheu, on a 1:1:1 mix of sand, peat and compost (Fig. S1C), with moisture and temperature depending on the local climate and natural rain only (Fig. S3). In autumn, this stubble was placed in experimental plots of winter oilseed rape and further incubated under the plants (Fig. S1D). When pseudothecia had appeared and finished to release spores, each stubble was washed and stored dry. The climate in the area is oceanic, and meteorological data were obtained from the INRA CLIMATIK database, for Le Rheu weather station, on an hourly basis. Cumulative temperature (Fig. S3A), rainfall (Fig. S3B) and days favourable for pseudothecial maturation were calculated (Fig. S3C). A day was considered favourable if the mean temperature was between 2 and 20°C and if the cumulative rainfall over the previous 11 days beforehand (including the day in question) exceeded 4 mm (Aubertot et al., 2006; Lô-Pelzer et al., 2009a). Given these parameter values, 64 favourable days were required for 50% of pseudothecia to reach maturation.

### 2.3 Image acquisition

Because we aimed at detecting black fruiting bodies on the dark and non homogeneous background of oilseed rape stubble, special care was taken with standardisation of pictures. To ease automated detection of stems, a picture of each group of 5 dry stems was taken on a blue background (PVC sheet Lastolite Colormatt electric blue). As mentioned before, BBQ sticks had been painted in blue. The barcoded label was always placed at the same place and included on the picture (Fig. S1EF). Attention was paid to have stems parallel to the small side of the picture, with crown end of the stems on the same side. To ensure the absence of stem shade on the picture, each group of 5 dry stems was placed on a glass plate, 16 cm above the blue PVC sheet. Two FotoQuantum LightPro 50 × 70cm softboxes were placed on both sides of the stubble with 4 daylight bulbs each (5400K, 30W). We checked that the bulbs were within the lower 45 angle to avoid reflection of the lights on the glass plate. Pictures were taken with a Nikon D5200 with an AF-S DX Micro Nikkor 40mm 1:2.8G lens, on a Kaiser Repro stand, with a wired remote control. Aperture was set at F22 for maximal depth of field, iso 125, daylight white balance. Pictures were saved as RGB images with a resolution of 6000 × 4000 pixels (Fig. S1F).

### 2.4 Image processing

#### 2.4.1 Pre-processing

For each digital image, the sample unique identification number (suid) was retrieved by reading the barcode using the ZBar library (Brown, 2018) and the stubble pieces were segmented using morphological image processing (Gonzalez and Woods, 2006). Then, each digital image was automatically split in several new images, each containing only one stem, and each being identified by their common suid and their stem number. Finally, each new image was cropped to focus on our region of interest, composed of only the 5 cm portion on crown end of the stem, i.e. where pseudothecia is mainly located.

#### 2.4.2 Data sets

To define our training and test sets, we selected 81 stems from pre-processed images to span the range of stubble origin, of canker severity classes, of pseudothecial density and of stubble colour. We included tricky cases like stubble with holes (pitch dark on the pictures) or moulded with other saprophytic fungi. Each of these images was manually processed by a single observer. Using GIMP 2.8 software (The GIMP team, 2018), a new binary picture was created where each of the pseudothecia seen was coloured. This provided us with pairs of files, one with the original picture; the other being an overlay containing the expert manually-detoured pseudothecia.

There was no expert manually-detoured stubble pieces because the problem of segmenting the foreground stubble from the blue background was negligible compared to segmenting the fruiting bodies from the stubble. Hence, each stubble was segmented using a classical unsupervised clustering-based thresholding method (Otsu, 1975).

Out of the 81 annotated images (i.e. ground truth), 20 were used as a training set for fruiting bodies detection by supervised learning (see 2.4.3) and 61 were used as a test set, to ensure the generalised trained method performed properly. To construct our sets, we randomly split our data and verified that the resulting fruiting body densities of each set were comparable, i.e. a mean density of 0.028 and 0.027 and a standard deviation of 0.030 and 0.023, respectively for training and test.

#### 2.4.3 Fruiting bodies detection

Following typical steps for machine-learning in image processing (LeCun et al., 2015; Sommer et al., 2011), we (a) derived several non-linear image features, using convolution operators (Gonzalez and Woods, 2006), to capture local image characteristics and (b) chose a state-of-the-art classifier to learn from the test set. We stated our problem as predicting the class of each pixel of each image as a function of the computed features, assuming that each individual pixel can be in either state F (fruiting bodies) or in state S (stubble or stem).

Concerning point (a), we computed classical features for colors (gaussian, spaces derived from RGB), edges (laplacian) and textures (eigenvalues of tensor structure and hessian). As a picture often contains informative characteristics at different level of organisation, we computed our features at different scales around every image’s pixel. Hence, for each feature, a maximum of 4 scales were computed with relevant range up to 7 pixels away. As for solving the classification problem, i.e. point (b), we chose to use gradient boosting (Friedman, 2001) as a recent and robust supervised ensemble learning method.

#### 2.4.4 Post-processing and quality assessment of the processing chain

We used complementary metrics, on the test set, to assess the quality of our model : (i) for each image, we computed the model accuracy i.e. the number of true positive and true negative pixels divided by the total amount of pixels (ii) and, for the whole test set, we computed the correlation between predicted and observed fraction of fruiting bodies pixels in states F (fruiting bodies) and S (stem).

As it is common in image processing, we decided to add some post-processing, which in this case aimed at improving even further the image processing chain including the detection routine. We chose a very simple, yet efficient post-process, consisting in a computer-assisted expert curation of the predicted fruiting bodies segmentation. An easy to use graphical user interface was developed for rapid assignation of a post-processed state, i.e. “correct” or “incorrect”, to each processed image by visually evaluating the predicted RGB image compared to the original one. Finally, we assessed the effects of year, field and severity before incubation on the probability of being classified as state “incorrect” by using a Generalised Linear Model (Binomial distribution and a logit link function) and Wald tests.

### 2.5 Statistical analysis of processed fruiting bodies data

We kept the post-processed images which were classified in state “correct” by the curator (n=2094) to assess the influence of (i) when and (ii) where the infected stems were collected, and (iii) the observed severity before incubation on the production of fruiting bodies. For each image *i* we considered the number of pixels in states F (fruiting bodies) and S (stem), i.e. *n_F,i_* and *n_S,i_*, and used a likelihood function based on 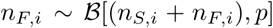 with a logit link function to build Generalised Linear Models and analyse the effects of year (4 levels), field (27 levels) and severity before incubation (6 levels) with Wald tests.

The whole image processing described in 2.4 was developed in Python using the scikit learn and image libraries (Pedregosa et al., 2011; van der Walt et al., 2014) while all the statistical analyses were performed with R (R Core Team, 2018).

## 3 Results

### 3.1 Fruiting bodies detection

As previously stated, our post-processing rely on an expert curation. On our test set, this process set aside 10 out of the 61 images (i.e. 16%). Over the curated set, this lead to a mean and median accuracy of 0.97 with a minimum at 0.87 and a maximum at 1. The adjusted R-squared between predicted and observed F/S is 0.87. For comparison, the adjusted R-squared without curation would have been 0.59. See also Figure 2 for representations of the model accuracies and predicted versus observed F/S.

**Figure 2:**
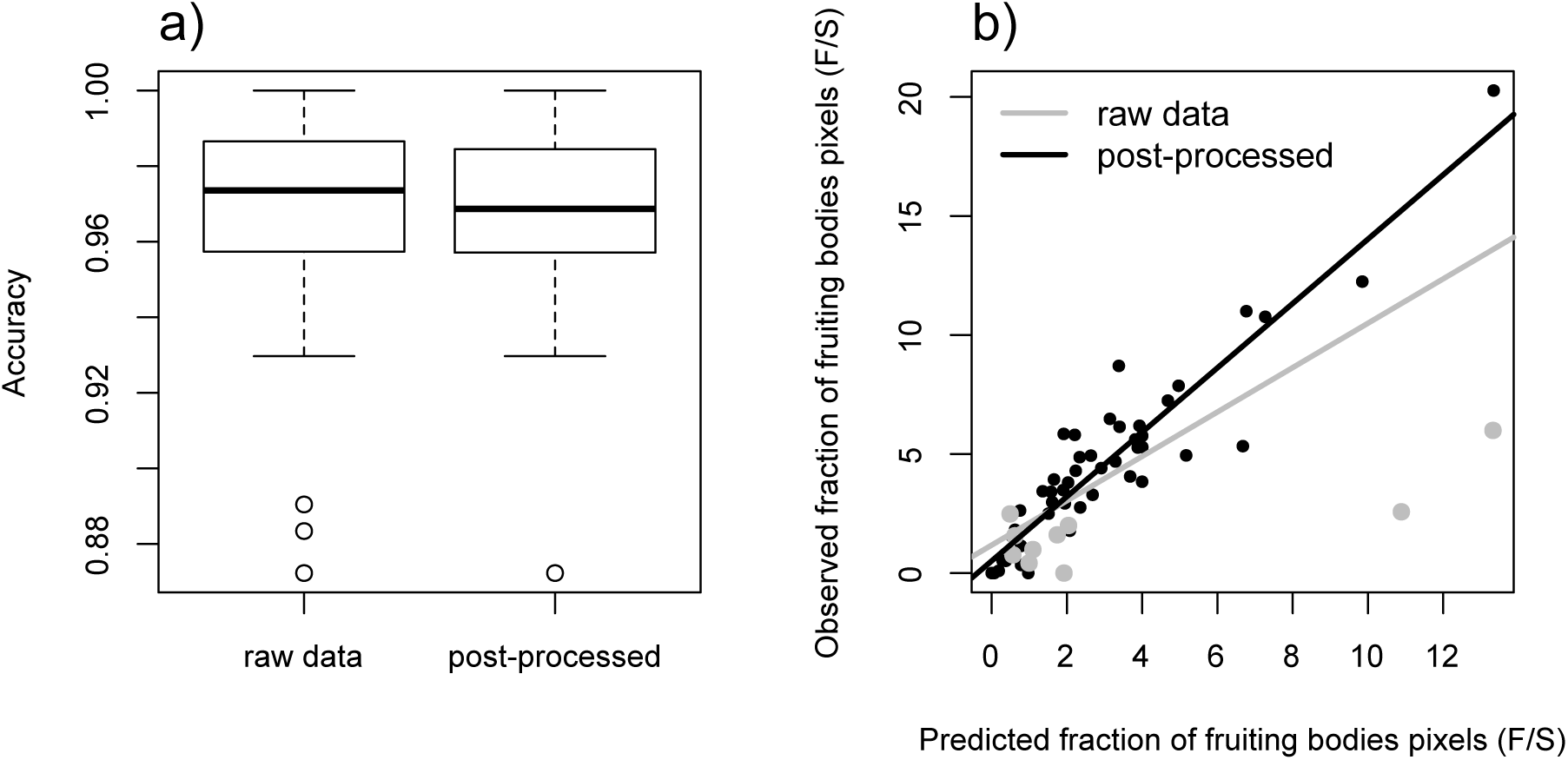
Fruiting bodies detection: a) model accuracies over the test set with or without curation, b) correlation between the predicted and observed F/S with (black) or without curation (gray).

**Figure 3:**
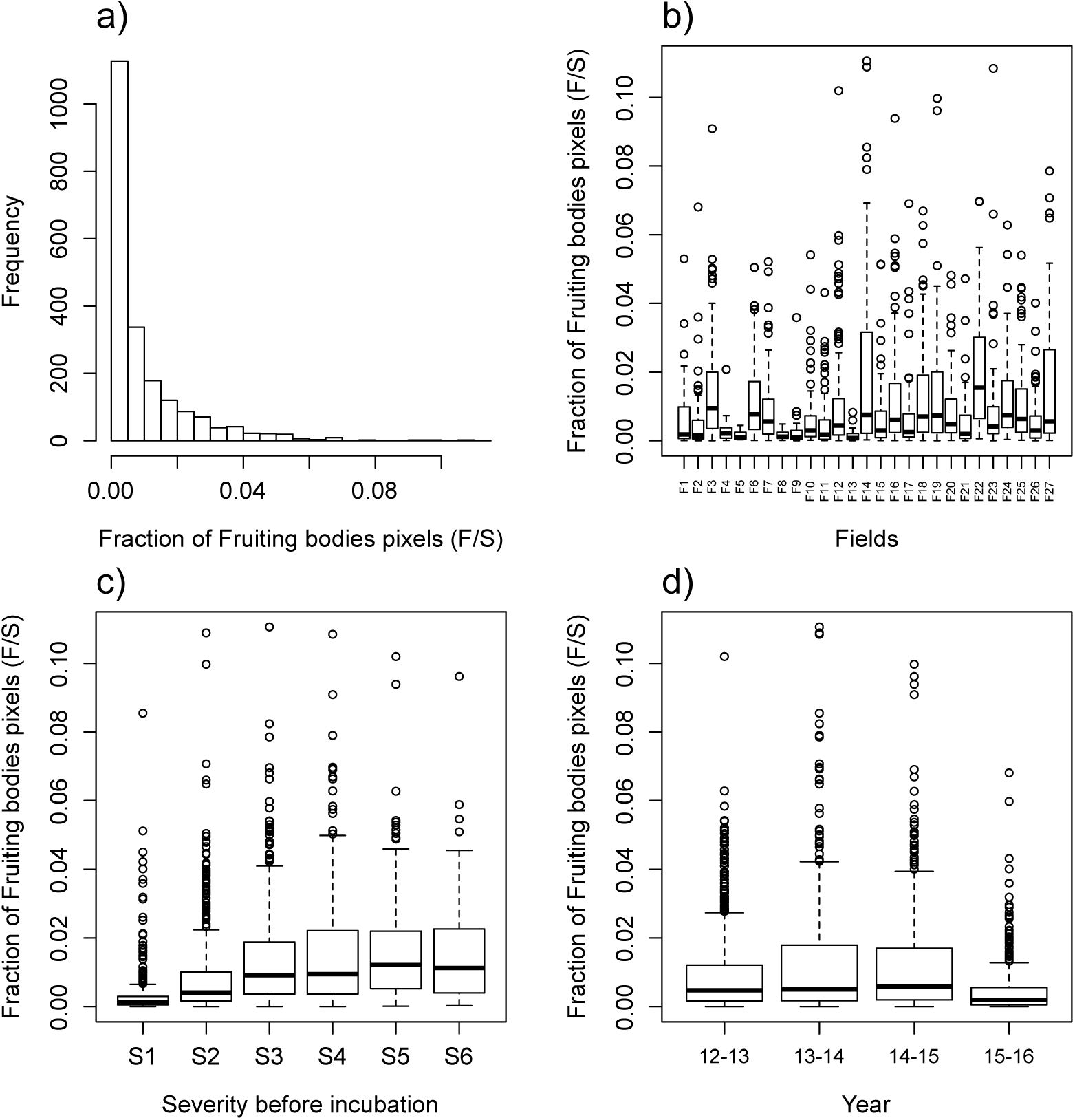
Graphical examination of the processed fraction of fruiting bodies on oilseed rape stem. a) shows the histogram of the all post-processed data set while b), c) and d) represent respectively how the fraction F/S change among the different fields (labelled F1 to F27), severity before incubation and year with boxplots.

Over the 2540 stems, 446 were classified as “incorrect” by the expert after post-processing. This percentage of discard (17.5%) is similar with what happened for the test set (16%). Examples of ‘correct’ and ‘incorrect’ segmentations are provided in figures 4 and 5. The year (*p* = 0.03) and field (*p* < 10^5^) variables had significant effects on the probability of being incorrectly segmented but only explained respectively 6.6% and 3.2% of the deviance while the effect of the severity before incubation of oilseed rape stems was not significant (*p* = 0.15). We explained these results by high level of humidity which occured near harvest in some years (i.e. 2014 and 2015) and fields hence induced a darkening of stems. Our segmentation method failed, by overestimating the level of fruiting bodies pixels, in these particular conditions.

**Figure 4:**
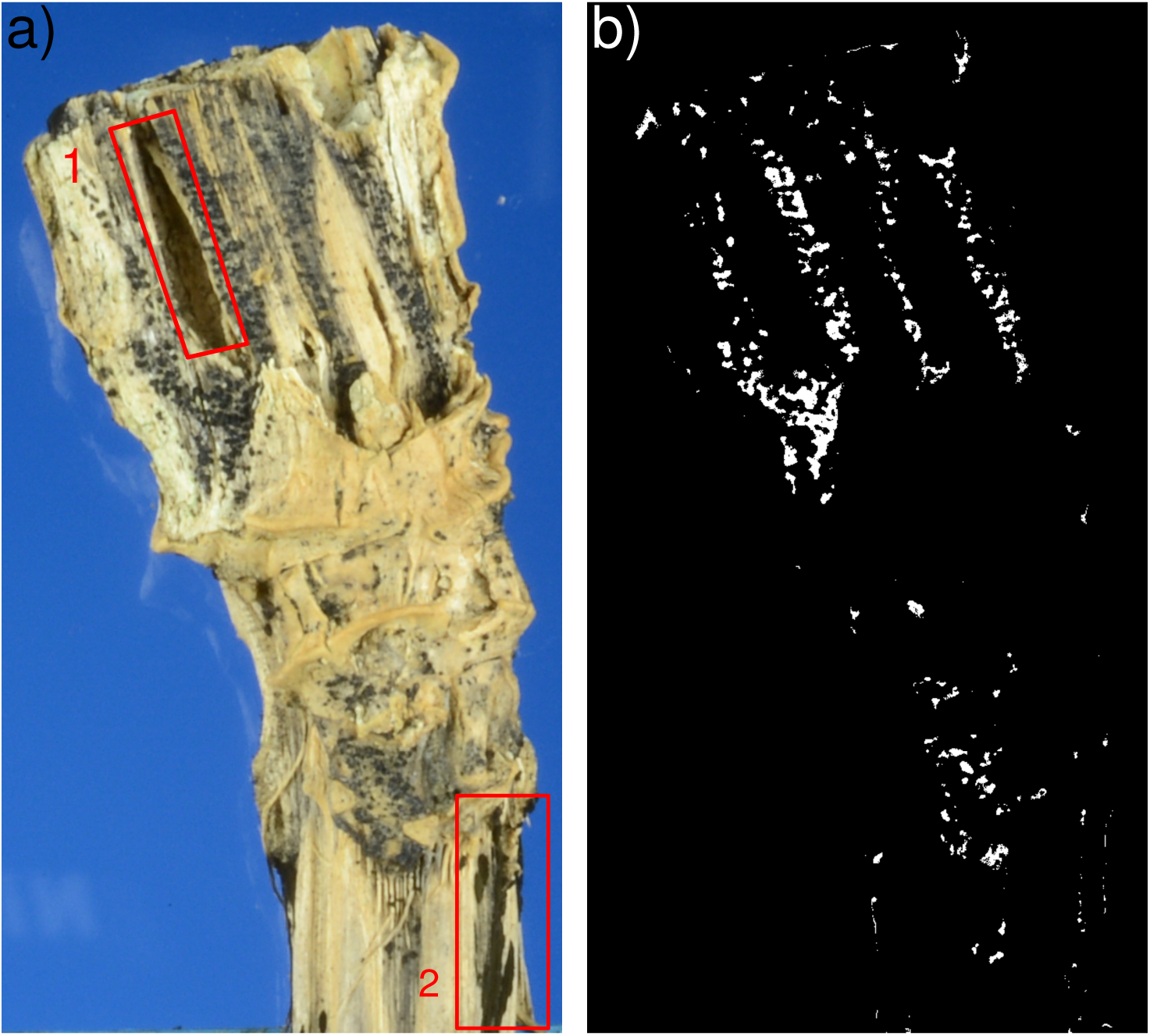
Illustrative example of a correct segmentation of fruiting bodies (b) (in white) from the RGB acquired image (a) despite the presence of hole (red rectangle 1) and other fungi (red rectangle 2) on the original oilseed rape stem after incubation.

**Figure 5:**
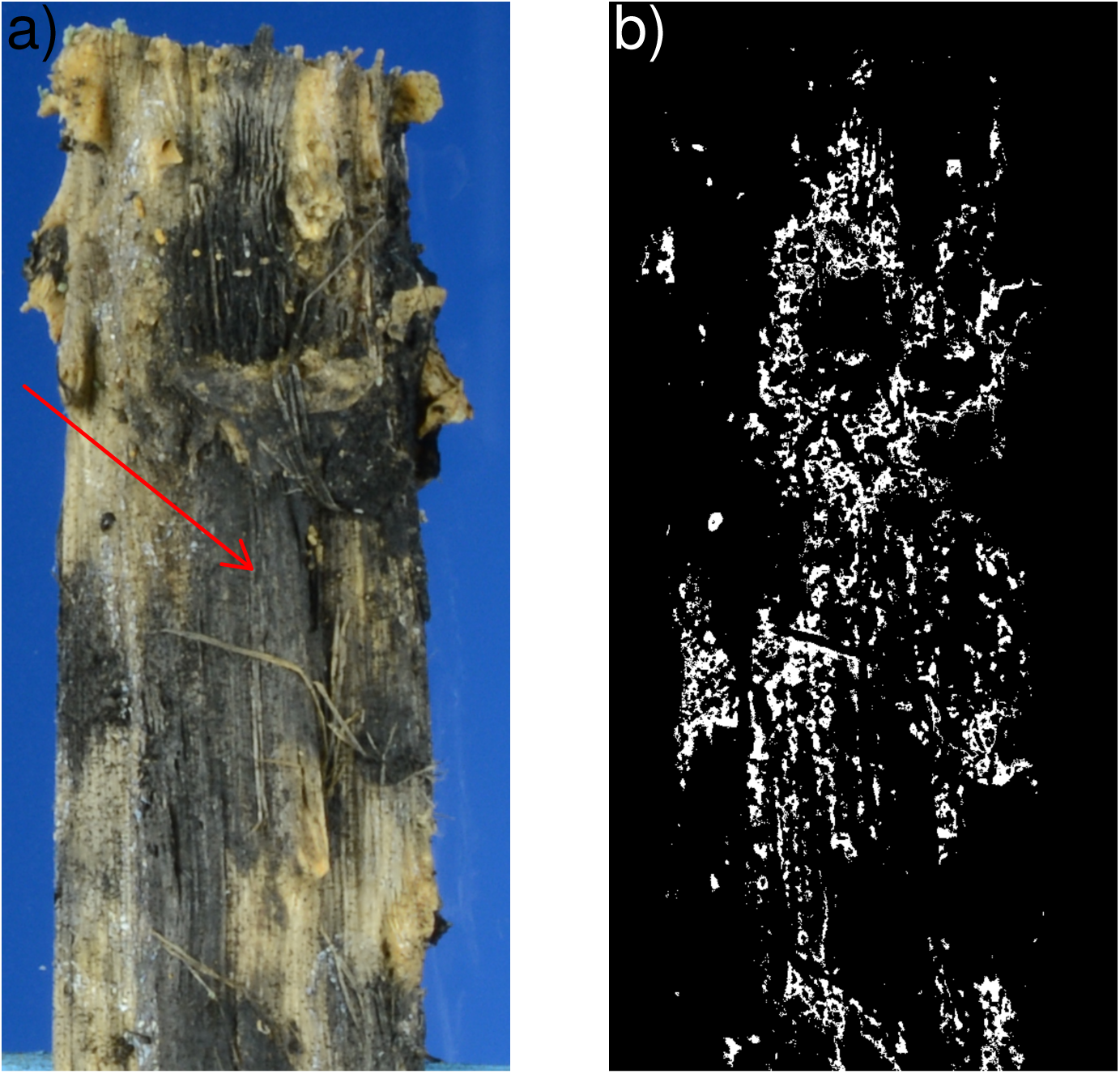
Illustrative example of an incorrect segmentation of fruiting bodies (b) (in white) from the RGB acquired image (a) where the overestimation of fruiting bodies was probably caused by the moisture-induced darkening of the oilseed rape stem (indicated by the red arrow on the RGB image).

### 3.2 Statistical analysis of processed fruiting bodies data

Our processing chain, whose components are illustrated in Figure 1, allowed us to quantify satisfactorily fruiting bodies on 2094 stems collected during 4 years in a oilseed rape production area. The distribution of the fraction of Fruiting bodies pixels (F/S) has a mode close to zero and exhibited a positive skewness with only a few outliers above 8%, which illustrates the low occurrence of fruiting bodies on oilseed rape stems (Fig. 3a). As suggested by graphical descriptive analysis (Figs. 3b-d), the statistical analyses confirmed the significant effects of the three categorical variables tested here (i.e. p-value < 10^−15^ for year, field and severity before incubation) on the presence of fruiting bodies on stems (Table S7). Year, field and severity before incubation explained respectively 6.4%, 15.9% and 22.1% of the deviance and these results were similar without the post-processing step (i.e. 8.5%, 16.3% and 19.4% explained by year, field and severity). The pairwise comparisons of least-square means pointed out a significant differences between all the considered fields, and, the 4 seasons during which stems were collected, 2015-2016 being the one with the lowest production of fruiting bodies and 2013-2014 the highest. The severity classes were logically ranked from the lowest to the highest except for classes S6 and S5 that were reversed (Table S9). This confirmed that severity at harvest is positively correlated with fruiting bodies production but suggested that class S6 (100%) should be merged with S5 (76-99%), at least from an epidemiological point of view. For supplementary information on these statistical analysis, see sections S-4 and S-5.

## 4 Discussion

High throughput phenotyping is a rich new ground upon which to base tomorrow’s epidemiological research. In this article, we achieved the construction of such a framework, binding automatic data extraction, using image processing, with statistical analysis of relevant explanatory variables. By careful curation and annotation of 81 numerical images of biological samples (i.e. ground truth), we managed to subsequently generate 2094 new reliable data. The gain in statistical power for testing the effects of covariates on pathogen development is obvious but, one can also note that this shifts the experimental bottleneck from time-consuming sample’s data extraction (low throughput) to experimental sampling *‘en masse’* (high throughput).

On top of its methological aspect, our current study is congruent with previous results on the blackleg of oilseed rape and opens the prospect to refine assessments. McGee and Emmett (1977) pointed to the fact that more pseudothecia appeared on stems with higher canker severity. However, they did not attempted to quantify fruiting bodies, and focused on liberated spores among three severity classes and four liberation dates. In their study, Marcroft et al. (2004b) visually estimated the percentage of area covered with pseudothecia and were able to process 320 stems to cover 2 years, 2 sites and 4 varieties. They did not find a significant effect of severity, but they had severity levels averaged over all stubble of a variety and not per individual stem. Lô-Pelzer et al. (2009b) followed individual stems and counted pseudothecia, insisting on the tediousness of the task on hundreds of stems. They were able to generate precise data from experimental plots and confirmed that the positive relationship between the number of pseudothecia and the severity of the blackleg also apply for farmers’ fields, though in the latter case more variation was observed. Thus, our results are in agreement with previous ones for canker severity classes 1 to 5, and suggest that new studies should be set up to understand discrepancies regarding class 6. Moreover, by developing an image-processing framework for quantifying pseudothecia on stubble we confirmed the statement of Lô-Pelzer et al. (2009b), who suggested that the use of digital images could be a way to improve the quantification of pseudothecia on stubble, and demonstrated that it enables one to increase the numbers of processed stems to thousands. Finally, our study is also in line with findings of Marcroft et al. (2004a) who identified a significant effect of host species and oilseed rape variety on both the visual density of pseudothecia and the numbers of liberated ascospores before suggesting that reduced potential for inoculum production could be a trait worth breeding for.

Our framework now provides a way to achieve sufficient throughput in data collection to study genotype effects more precisely than previous studies (e.g. (Lô-Pelzer et al., 2009b; Marcroft et al., 2004b)). Indeed, because Marcroft et al. (2004b) did not track canker severity on individual stems, the observed reduction of fruiting bodies jointly results from reduced canker severity, and potentially from the genotype. Lô-Pelzer et al. (2009b) were able to see the effect of a genotype with quantitative resistance on the distributions of severities, though they observed no effect of the genotype at a given severity. Screening more genotypes is needed to identify if there is variation for this trait in current germplasm, and our framework provides a way to do so. The number of produced pseudothecia is known to change with the year and the location of the sampling (Lô-Pelzer et al., 2009b). Our results are in agreement with these findings, although in our study different host varieties might further contribute to amplify these effects. From the biotrophic and asymptomatic presence of the fungus in the stem, visible cankers appear progressively when crop matures. Moreover, the delay between infection and symptom appearance (i.e. incubation period) is known to vary between host-plants in a population and the distribution of the incubation period could also change with some covariates like the host-age when the infection occurs (Leclerc et al., 2014). Then, when collecting disease data for a multi-year and site study, the sampling can occur at different stages in these processes, and thus introduce more variability in the relationship between the visual canker severity and the resulting fruiting bodies produced. We believe that our framework which allows the quantification of pseudothecia on a large number of stubble could be used to study precisely the dynamics of pseudothecial appearance and investigate how it may be influenced by year, field and potentially cropping practice effects. It has been demonstrated that during pseudothecial maturation, ascospores appear at a rate following a sigmoid-shaped function (Wherrett et al., 2004) that is affected by climate (Naseri et al., 2009); chemical treatment (Wherrett et al., 2003) or cropping practice after harvest (McCredden et al., 2018). In combination with methods for ascospores counting, the use of image-based quantification of pseudothecia could help to disentangle pseudothecial maturation and ascospores emission by providing large and precise data-sets. Then, one would be able to estimate the distributions of the i) time to ascospores emission (pseudothecial maturation), ii) the number of spores produced by pseudothecium and iii) the emission function that are key processes for understanding and predicting the initiation of *L. maculans* epidemics in oilseed rape crops.

Yet, we must keep in mind that our current (and futur) results depends heavily on the quality of the imaging framework for data extraction, which involve many different processes. For example, in fig. 2, one can note that, in our study, the median accuracy is slightly worsen by the post-process expert curation while the adjusted R-squared is extremely improved. It highlights the usefulness of processed-images curation by an expert as the human operator can consider image features that were not not captured by the pixel-wise accuracy metric to filter the predictions. One way to remove, or at least lessen, the need for human post-processing could be to enforce object detection instead of pixel-wise classification (Sommer et al., 2011; Ren et al., 2017). An even more advanced use of imaging would be to use new imaging sensors (e.g. hyperspectral, chlorophyll fluorescence) in combination with machine learning methods (e.g. deep learning) which are already used in plant phenotyping (Barr et al., 2017; Pound et al., 2016) and expanding in phytopathology (Moghadam et al., 2017; Wang et al., 2017). Hence, our whole framework could benefit from a cascade detection (Zhou, 2012) of fruiting bodies with different steps involving different sensors and/or algorithms. For example, a first step could be to apply our current algorithm on RGB images then, if image acquisition, data extraction and expert curation was prompt enough, we could acquire new images of expert rejected stubbles with advanced imaging sensors (e.g. hyperspectral) hoping that this type of signal lead to improved accuracy over RGB color space. Such a combination of sensors could increase the overall accuracy while still cutting costs (as opposed to scanning all samples with any and every sensors available).

Besides improving the image processing, we could also improve the epidemiological models for predicting disease development. For instance, we may take advantage of the high-throughput epidemiological data stream by developing predictive risk models using state-of-the-art machine learning incorporating readily available covariates (e.g. climate, remote sensed land-use, …). On the mechanistic side of things, it would be also interesting to use our phenotyping framework to produce time-space data at the landscape scale and feed previously developed spatio-temporal models (Bousset et al., 2015) and refine the challenging quantification of epidemiological processes such as anisotropic dispersal between locations, between-year inoculum transmission and the susceptibility of host-fields. Finally, the combination of high-throughput image-based epidemic data with current next generation sequencing methods appears as an interesting perspective to tackle demo-genetics questions to understand better of the genetic structures of host-plants influence the population genetics of *L. maculans* among a cultivated area and improve the management of plant resistances for a durable control of the disease (Bousset et al., 2018).

## Acknowledgements

We thank Farmers near Le Rheu who allowed us to assess disease in their fields. We thank Yannick Lucas, Hervé Picault, Magali Ermel, Claude Domin and Ronan Le Cointe for technical assistance, and Sylvain Prigent from the Biogenouest plateform for his advice on image analysis. We are gratefull to the INRA CLIMATIK database for the weather data. This work benefited from the financial support of INRA the French National Institute for Agronomical Research, and from ANR the French National Research Agency programs Agriculture et Dveloppement Durable grant ANR-05-PADD-05, CEDRE, AGROBIOSPHERE grant ANR-11-AGRO-003-01. The authors declare the absence of conflict of interest. MP and NP developed the image processing code, LB carried out the case study experiment, ML and NP performed statistical analyses. LB, NP and ML conceived and designed the study and prepared the manuscript, read and approved by all authors. Source code is available upon request.

## Supplementary materials

### S-1 Image acquisition process

**Figure S-1:**
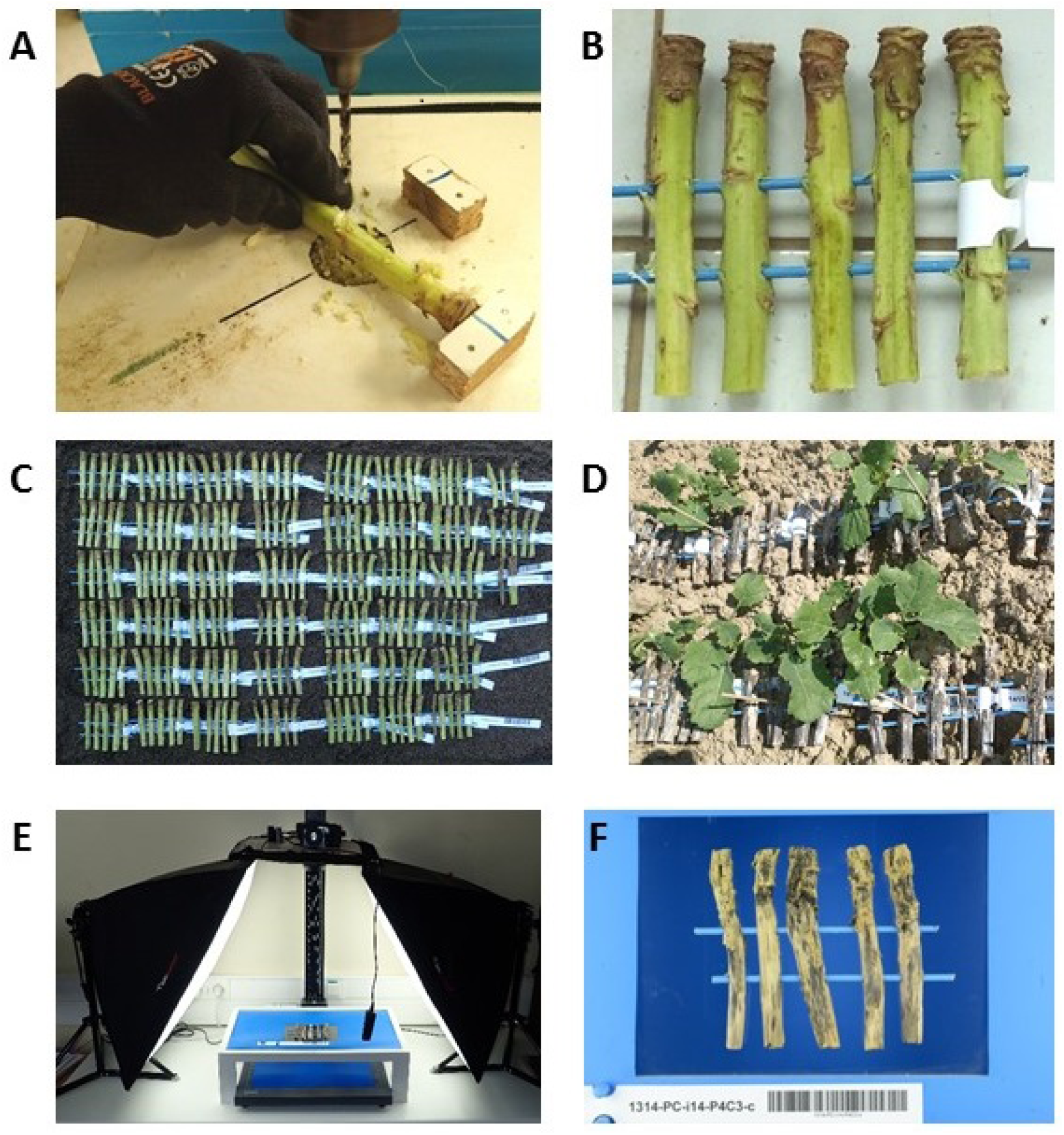
Illustrations of the process. A. Two 5 mm diameter holes were drilled in 10 cm long stems cut at crown for canker severity assessment. B. Keeping track of disease severity, stems were grouped by 5 on two wooden BBQ sticks painted in blue and labelled with a barcode. C. Stubble was matured outside over summer. D. Stubble was transferred in experimental oilseed rape field to further mature over autumn and winter. E. Standardized pictures of the washed and dried stubble were taken on a glass plate over blue background with daylight bulbs. F. Resulting standardized files contained the barcoded label and dry stubble.

### S-2 Image processing chain

**Figure S-2:**
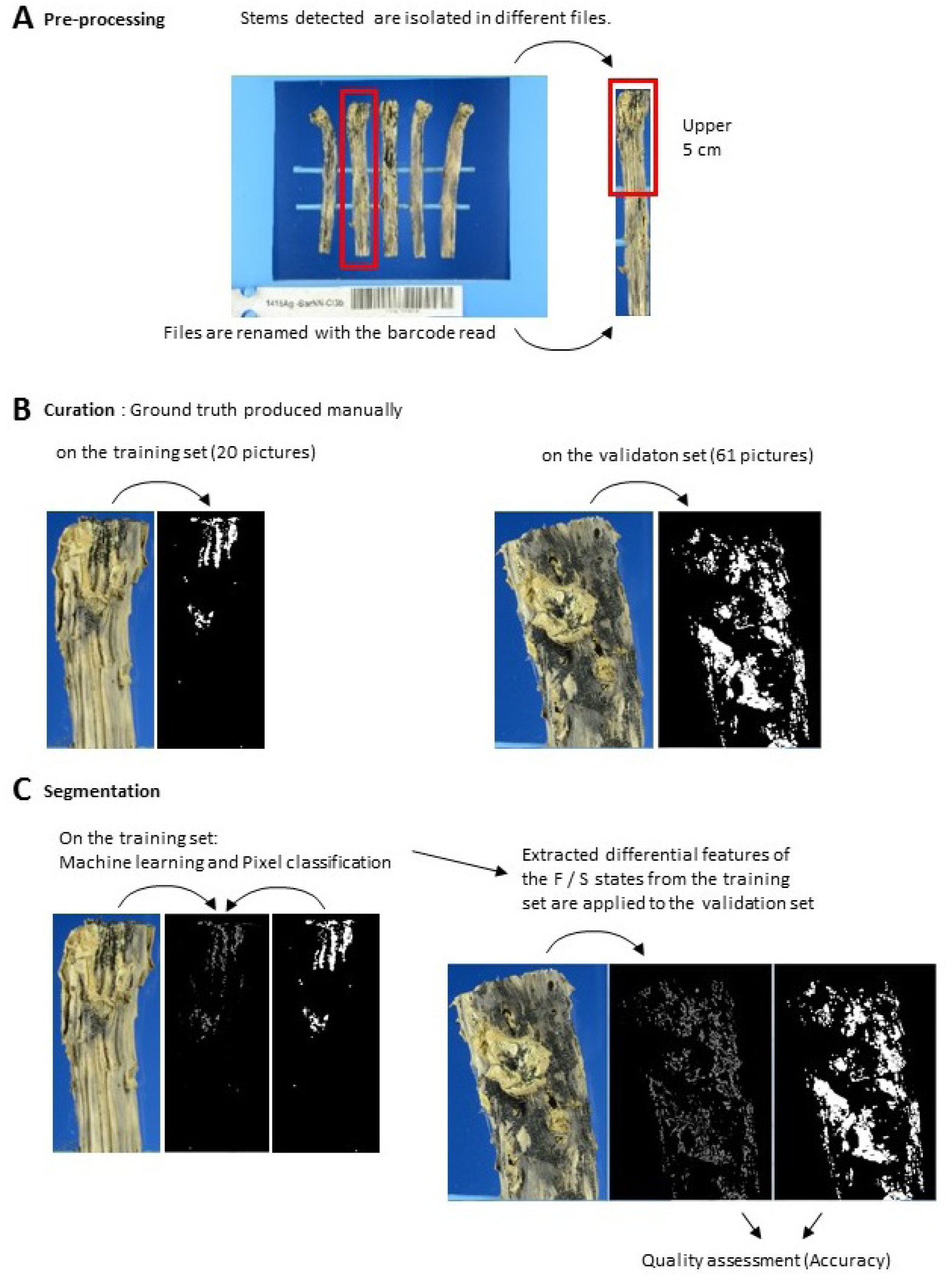
Illustrations of the picture processing chain. A. Pre-processing includes the segmentation of individual stems, isolated into separate new files, renamed with the barecode read, and cropped to the upper 5 cm. B. Curation includes the manual segmentation by an expert, both for the training set (20 pictures) and the validation set (61 pictures). C. Segmentation by machine learning on the training set followed by pixel classification according to ground truth enables to extract features differential between pixels in the state S (stem) or F(fruiting bodies), respectively. These differential features are then applied to the validation set. Quality assessment is performed by comparing the segmented pictures with the corresponding ground truth, calculating accuracy.

### S-3 Assessment of climatic conditions during pseudothecial maturation

**Figure S-3:**
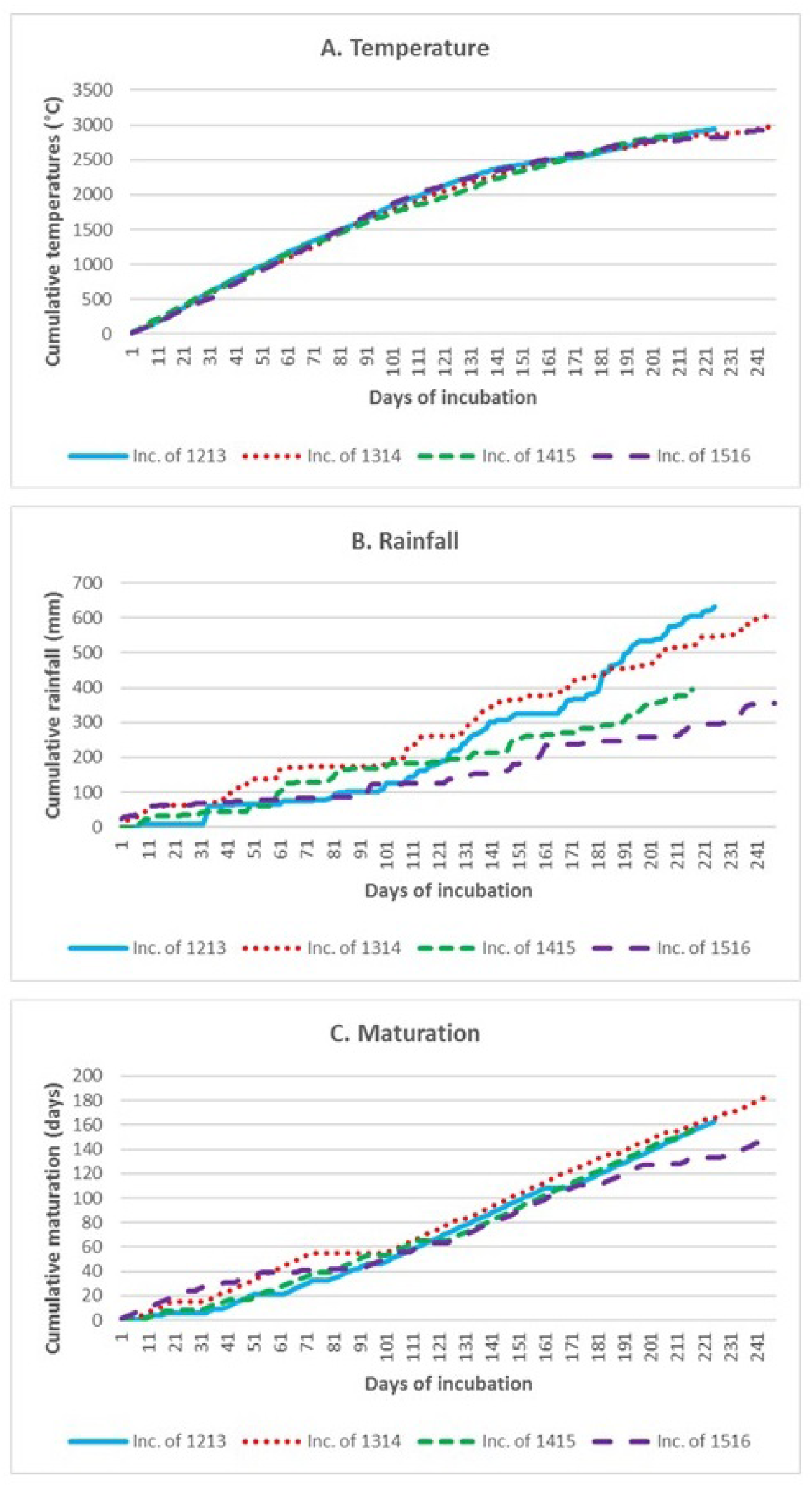
Weather and pseudothecial maturation data during the incubation of stubble of four cropping seasons 2012-2013 (Inc. of 1213) to 2015-2016 (Inc. of 1516. A. Mean daily temperature was cumulated over the course of each experiment. B. Cumulative rainfall. C. Cumulative numbers of days favourable for the maturation of pseudothecia. A day was considered favourable if the mean temperature was between 2 and 20C and if the cumulative rainfall over the previous 11 days beforehand (including the day in question) exceeded 4 mm (Aubertot et al. 2006; L-Pelzer et al. 2009). Given these parameter values, 64 favourable days are required for 50% of pseudothecia to reach maturation. Meteorological data were obtained from the INRA CLIMATIK database, for Le Rheu weather station, on an hourly basis.

### S-4 Supplementary information on post-processing

#### S-4.1 Description of the data ranked as correct (grey)or incorrect (black) depending on a) field; b) year; c) disease severity. d) ranking depending on the proportion of fruiting bodies pixels

**Figure S-4:**
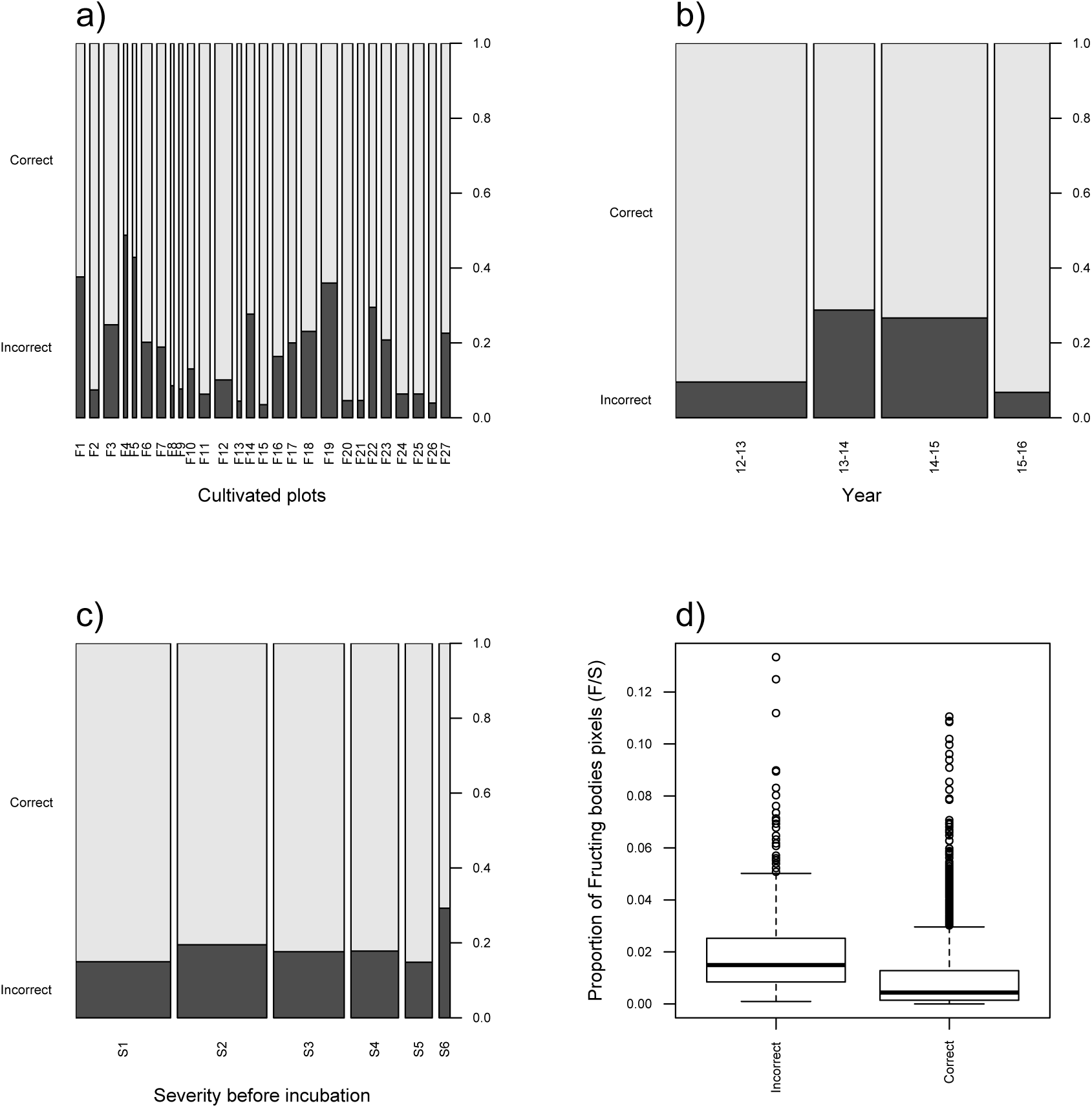
Graphical examination of the post-processed data.

**Table S-1:** Summary statistics of post-processed data and related covariates

| Year | Field | Severity | Number of pixels S | Number of pixels F | $F/S \times 100$ | Post-processed state |
| --- | --- | --- | --- | --- | --- | --- |
| 2012-2013:943 | F12 : 178 | S1:708 | Min. :141760 | Min. : 0 | Min. : 0.0000 | Incorrect: 446 |
| 2013-2014:438 | F19 : 164 | S2:667 | 1st Qu.:260122 | 1st Qu.: 530 | 1st Qu.: 0.1875 | Correct :2094 |
| 2014-2015:761 | F18 : 156 | S3:527 | Median :297408 | Median : 1756 | Median : 0.5900 |  |
| 2015-2016:398 | F3 : 153 | S4:354 | Mean :302215 | Mean : 3662 | Mean : 1.1700 |  |
|  | F24 : 126 | S5:202 | 3rd Qu.:337670 | 3rd Qu.: 4798 | 3rd Qu.: 1.5800 |  |
|  | F11 : 111 | S6: 82 | Max. :577404 | Max. :69123 | Max. :13.3400 |  |
|  | (Other):1652 |  |  |  |  |  |

#### S-4.2 Statistical analysis

**Table S-2:** Analysis of deviance and Wald-tests for the analysis of post-processed data

| Variable | Df | p-value | Explained deviance |
| --- | --- | --- | --- |
| Year | 3 | 0.03 | 6.6 % |
| Plot | 24 | $< 6 \times 10^6$ | 3.2 % |
| Severity | 5 | 0.15 | 0.5 % |

### S-5 Supplementary information on the analysis of fruiting bodies data

#### S-5.1 Description of post-processed data

**Table S-3:**
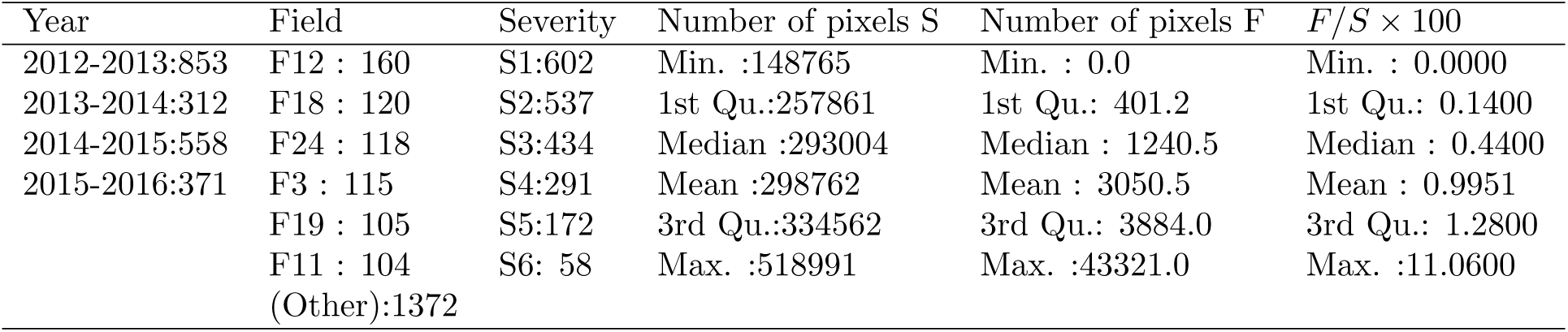
Summary statistics of processed data and related covariates

| Year | Field | Severity | Number of pixels S | Number of pixels F | $F/S \times 100$ |
| --- | --- | --- | --- | --- | --- |
| 2012-2013:853 | F12 : 160 | S1:602 | Min. :148765 | Min. : 0.0 | Min. : 0.0000 |
| 2013-2014:312 | F18 : 120 | S2:537 | 1st Qu.:257861 | 1st Qu.: 401.2 | 1st Qu.: 0.1400 |
| 2014-2015:558 | F24 : 118 | S3:434 | Median :293004 | Median : 1240.5 | Median : 0.4400 |
| 2015-2016:371 | F3 : 115 | S4:291 | Mean :298762 | Mean : 3050.5 | Mean : 0.9951 |
|  | F19 : 105 | S5:172 | 3rd Qu.:334562 | 3rd Qu.: 3884.0 | 3rd Qu.: 1.2800 |
|  | F11 : 104 | S6: 58 | Max. :518991 | Max. :43321.0 | Max. :11.0600 |
|  | (Other):1372 |  |  |  |  |

**Table S-4:** Contingency table for variables Field and Year

|  | 2012-2013 | 2013-2014 | 2014-2015 | 2015-2016 |
| --- | --- | --- | --- | --- |
| F1 | 0 | 0 | 93 | 0 |
| F2 | 0 | 0 | 0 | 94 |
| F3 | 0 | 0 | 153 | 0 |
| F4 | 0 | 41 | 0 | 0 |
| F5 | 0 | 42 | 0 | 0 |
| F6 | 109 | 0 | 0 | 0 |
| F7 | 90 | 0 | 0 | 0 |
| F8 | 35 | 0 | 0 | 0 |
| F9 | 0 | 0 | 0 | 39 |
| F10 | 69 | 0 | 0 | 0 |
| F11 | 40 | 0 | 0 | 71 |
| F12 | 105 | 0 | 0 | 73 |
| F13 | 0 | 0 | 0 | 45 |
| F14 | 0 | 83 | 0 | 0 |
| F15 | 85 | 0 | 0 | 0 |
| F16 | 0 | 0 | 110 | 0 |
| F17 | 0 | 0 | 85 | 0 |
| F18 | 0 | 0 | 156 | 0 |
| F19 | 0 | 0 | 164 | 0 |
| F20 | 109 | 0 | 0 | 0 |
| F21 | 65 | 0 | 0 | 0 |
| F22 | 0 | 78 | 0 | 0 |
| F23 | 0 | 101 | 0 | 0 |
| F24 | 126 | 0 | 0 | 0 |
| F25 | 110 | 0 | 0 | 0 |
| F26 | 0 | 0 | 0 | 76 |
| F27 | 0 | 93 | 0 | 0 |

**Table S-5:**
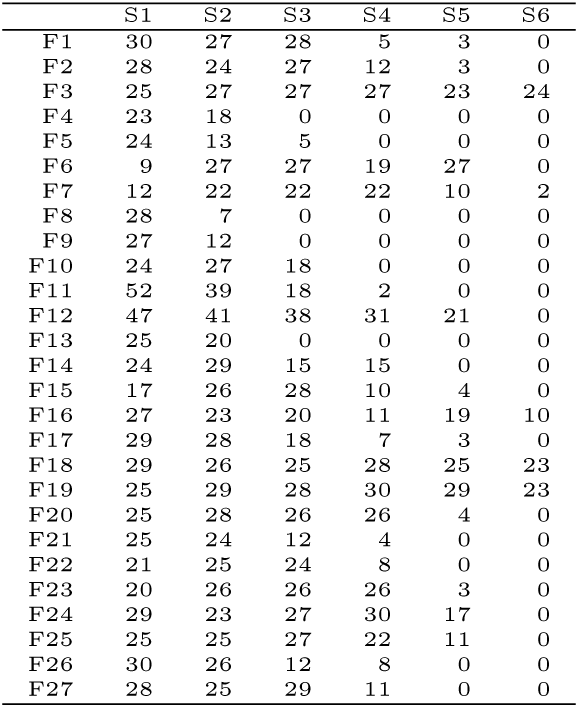
Contingency table for variables Field and Severity before incubation

|  | S1 | S2 | S3 | S4 | S5 | S6 |
| --- | --- | --- | --- | --- | --- | --- |
| F1 | 30 | 27 | 28 | 5 | 3 | 0 |
| F2 | 28 | 24 | 27 | 12 | 3 | 0 |
| F3 | 25 | 27 | 27 | 27 | 23 | 24 |
| F4 | 23 | 18 | 0 | 0 | 0 | 0 |
| F5 | 24 | 13 | 5 | 0 | 0 | 0 |
| F6 | 9 | 27 | 27 | 19 | 27 | 0 |
| F7 | 12 | 22 | 22 | 22 | 10 | 2 |
| F8 | 28 | 7 | 0 | 0 | 0 | 0 |
| F9 | 27 | 12 | 0 | 0 | 0 | 0 |
| F10 | 24 | 27 | 18 | 0 | 0 | 0 |
| F11 | 52 | 39 | 18 | 2 | 0 | 0 |
| F12 | 47 | 41 | 38 | 31 | 21 | 0 |
| F13 | 25 | 20 | 0 | 0 | 0 | 0 |
| F14 | 24 | 29 | 15 | 15 | 0 | 0 |
| F15 | 17 | 26 | 28 | 10 | 4 | 0 |
| F16 | 27 | 23 | 20 | 11 | 19 | 10 |
| F17 | 29 | 28 | 18 | 7 | 3 | 0 |
| F18 | 29 | 26 | 25 | 28 | 25 | 23 |
| F19 | 25 | 29 | 28 | 30 | 29 | 23 |
| F20 | 25 | 28 | 26 | 26 | 4 | 0 |
| F21 | 25 | 24 | 12 | 4 | 0 | 0 |
| F22 | 21 | 25 | 24 | 8 | 0 | 0 |
| F23 | 20 | 26 | 26 | 26 | 3 | 0 |
| F24 | 29 | 23 | 27 | 30 | 17 | 0 |
| F25 | 25 | 25 | 27 | 22 | 11 | 0 |
| F26 | 30 | 26 | 12 | 8 | 0 | 0 |
| F27 | 28 | 25 | 29 | 11 | 0 | 0 |

**Table S-6:** Contingency table for variables Year and Severity before incubation

|  | S1 | S2 | S3 | S4 | S5 | S6 |
| --- | --- | --- | --- | --- | --- | --- |
| 2012-2013 | 242 | 239 | 210 | 160 | 90 | 2 |
| 2013-2014 | 140 | 136 | 99 | 60 | 3 | 0 |
| 2014-2015 | 165 | 160 | 146 | 108 | 102 | 80 |
| 2015-2016 | 161 | 132 | 72 | 26 | 7 | 0 |

#### S-5.2 Statistical analyses

**Table S-7:** Analysis of deviance and Wald-tests for the analysis of post-processed data (white) and non post-processed data (grey).

| Variable | Df | p-value | Explained deviance |
| --- | --- | --- | --- |
| Year | 3 | $< 2.2 \times 10^{-16}$ | 6.4 % |
| Field | 26 | $< 2.2 \times 10^{-16}$ | 15.9 % |
| Severity | 5 | $< 2.2 \times 10^{-16}$ | 22.1 % |
| Year | 3 | $< 2.2 \times 10^{-16}$ | 8.5 % |
| Field | 26 | $< 2.2 \times 10^{-16}$ | 16.3 % |
| Severity | 5 | $< 2.2 \times 10^{-16}$ | 19.4 % |

**Table S-8:** Pairwise comparisons of least-square means for the explanatory variable Year (R output)

| Year | prob | SE | asympt.LCL | asympt.UCL | Group |
| --- | --- | --- | --- | --- | --- |
| 2015-2016 | 0.0050 | 6.9673e-06 | 0.0050 | 0.0050 | 1 |
| 2012-2013 | 0.0094 | 5.9778e-06 | 0.0094 | 0.0094 | 2 |
| 2014-2015 | 0.0122 | 8.5705e-06 | 0.0122 | 0.0122 | 3 |
| 2013-2014 | 0.0145 | 1.2055e-05 | 0.0144 | 0.0145 | 4 |

**Table S-9:**
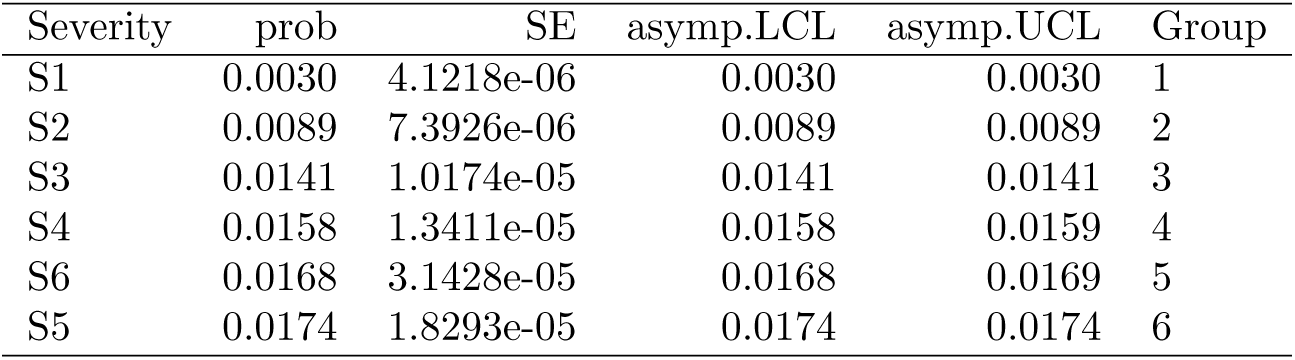
Pairwise comparisons of least-square means for the explanatory variable Severity before incubation (R output)

| Severity | prob | SE | asympt.LCL | asympt.UCL | Group |
| --- | --- | --- | --- | --- | --- |
| S1 | 0.0030 | 4.1218e-06 | 0.0030 | 0.0030 | 1 |
| S2 | 0.0089 | 7.3926e-06 | 0.0089 | 0.0089 | 2 |
| S3 | 0.0141 | 1.0174e-05 | 0.0141 | 0.0141 | 3 |
| S4 | 0.0158 | 1.3411e-05 | 0.0158 | 0.0159 | 4 |
| S6 | 0.0168 | 3.1428e-05 | 0.0168 | 0.0169 | 5 |
| S5 | 0.0174 | 1.8293e-05 | 0.0174 | 0.0174 | 6 |

**Table S-10:**
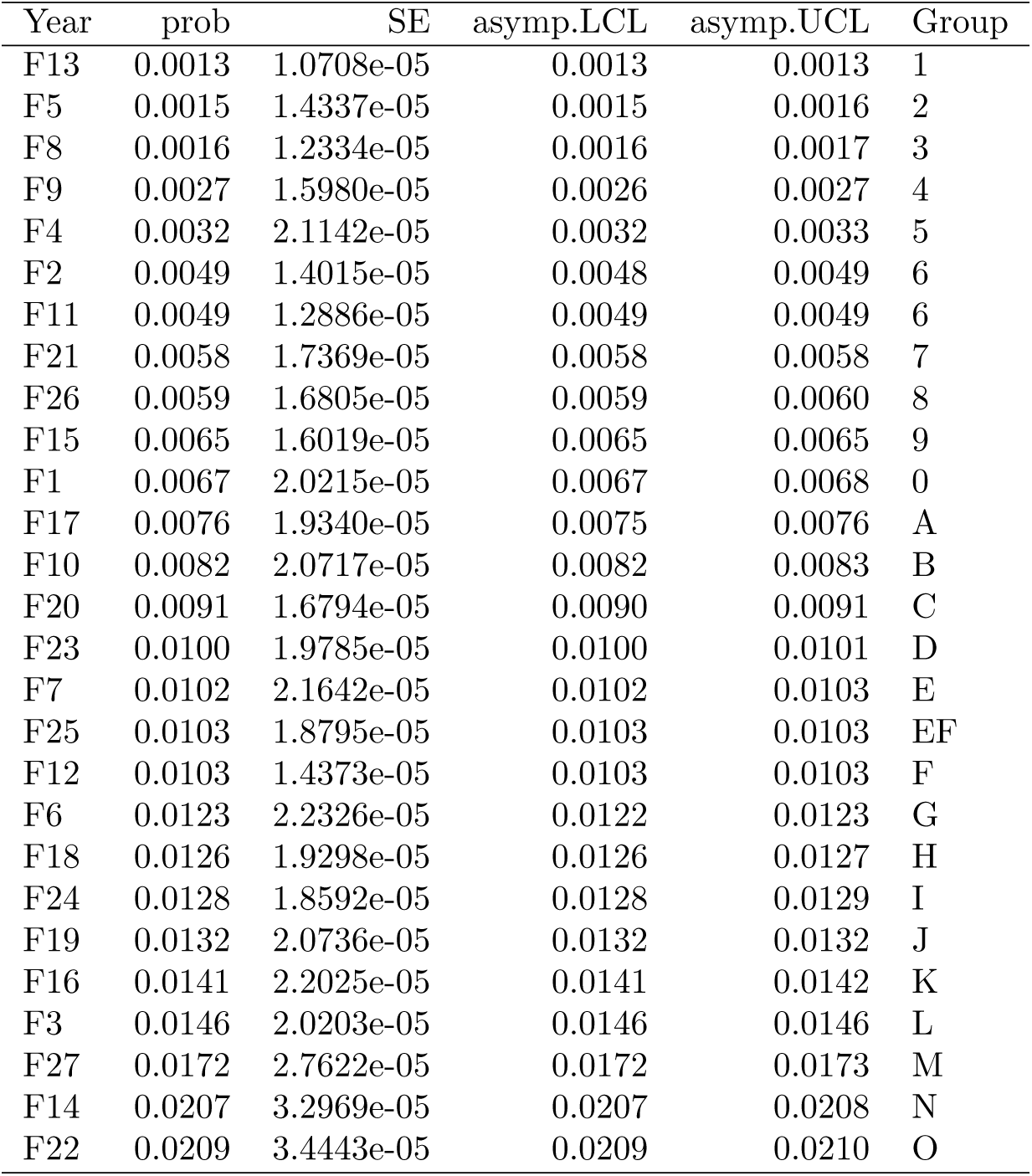
Pairwise comparisons of least-square means for the explanatory variable Field (R output)

## References

Aubertot, J., Salam, M. U., Diggle, A. J., Dakowska, S., and Jedryczka, M. (2006). Simmat, a new dynamic module of blackleg sporacle for the prediction of pseudothecia maturation of l. maculans/l. biglobosa species complex. parameterisation and evaluation in polish conditions. IOBC WPRS BULLETIN, 29(7):277.

Bailey, D., Kleczkowski, A., and Gilligan, C. (2004). Epidemiological dynamics and the efficiency of biological control of soil-borne disease during consecutive epidemics in a controlled environment. New Phytologist, 161(2):569–575.

Barr, P., Stver, B. C., Mller, K. F., and Steinhage, V. (2017). LeafNet: A computer vision system for automatic plant species identification. Ecological Informatics, 40: 50–56.

Bousset, L. and Chèvre, A.-M. (2013). Stable epidemic control in crops based on evolutionary principles: adjusting the metapopulation concept to agro-ecosystems. Agriculture, ecosystems & environment, 165: 118–129.

Bousset, L., Jumel, S., Garreta, V., Picault, H., and Soubeyrand, S. (2015). Transmission of leptosphaeria maculans from a cropping season to the following one. Annals of Applied Biology, 166(3):530–543.

Bousset, L., Jumel, S., Picault, H., Domin, C., Lebreton, L., Ribulé, A., and Delourme, R. (2016). An easy, rapid and accurate method to quantify plant disease severity: application to phoma stem canker leaf spots. European Journal of Plant Pathology, 145(3):697–709.

Bousset, L., Sprague, S. J., Thrall, P. H., and Barrett, L. G. (2018). Spatio-temporal connectivity and host resistance influence evolutionary and epidemiological dynamics of the canola pathogen leptosphaeria maculans. Evolutionary Applications.

Brady, C., Noll, L., Saleh, A., and Little, C. (2011). Disease severity and microsclerotium properties of the sorghum sooty stripe pathogen, ramulispora sorghi. Plant disease, 95(7):853–859.

Brown, J. (2018). Zbar bar code reader. Available at zbar.sourceforge.net.

Burt, P. J., Rosenberg, L., Rutter, J., Ramirez, F., and Gonzales O, H. (1999). Forecasting the airborne spread of mycosphaerella fijiensis, a cause of black sigatoka disease on banana: estimations of numbers of perithecia and ascospores. Annals of applied biology, 135(1):369–377.

Camargo, A. and Smith, J. (2009). An image-processing based algorithm to automatically identify plant disease visual symptoms. Biosystems Engineering, 102(1):9–21.

Delourme, R., Bousset, L., Ermel, M., Duffe, P., Besnard, A.-L., Marquer, B., Fudal, I., Linglin, J., Chadoeuf, J., and Brun, H. (2014). Quantitative resistance affects the speed of frequency increase but not the diversity of the virulence alleles overcoming a major resistance gene to leptosphaeria maculans in oilseed rape. Infection, genetics and evolution, 27: 490–499.

Friedman, J. H. (2001). Greedy Function Approximation: A Gradient Boosting Machine. The Annals of Statistics, 29(5):1189–1232.

Gilligan, C. A. (2002). An epidemiological framework for disease management. Advances in botanical research, 38(1):64.

Gonzalez, R. C. and Woods, R. E. (2006). Digital Image Processing (3rd Edition). Prentice-Hall, Inc., Upper Saddle River, NJ, USA.

Hamelin, F. M., Castel, M., Poggi, S., Andrivon, D., and Mailleret, L. (2011). Seasonality and the evolutionary divergence of plant parasites. Ecology, 92(12):2159–2166.

Karisto, P., Hund, A., Yu, K., Anderegg, J., Walter, A., Mascher, F., McDonald, B. A., and Mikaberidze, A. (2018). Ranking quantitative resistance to septoria tritici blotch in elite wheat cultivars using automated image analysis. Phytopathology, 108(5):568–581.

Leclerc, M., Dor, T., Gilligan, C. A., Lucas, P., and Filipe, J. A. N. (2014). Estimating the delay between host infection and disease (incubation period) and assessing its significance to the epidemiology of plant diseases. PLOS ONE, 9(1):1–15.

LeCun, Y., Bengio, Y., and Hinton, G. (2015). Deep learning. Nature, 521(7553):436–444.

Lô-Pelzer, E., Aubertot, J.-N., Bousset, L., Pinochet, X., and Jeuffroy, M.-H. (2009a). Phoma stem canker (leptosphaeria maculans/l. biglobosa) of oilseed rape (brassica napus): is the g2 disease index a good indicator of the distribution of observed canker severities? European journal of plant pathology, 125(4):515–522.

Lô-Pelzer, E., Aubertot, J.-N., David, O., Jeuffroy, M.-H., and Bousset, L. (2009b). Relationship between severity of blackleg (leptosphaeria maculans/l. biglobosa species complex) and subsequent primary inoculum production on oilseed rape stubble. Plant Pathology, 58(1):61–70.

Mahlein, A.-K. (2016). Plant disease detection by imaging sensors-parallels and specific demands for precision agriculture and plant phenotyping. Plant Disease, 100(2):241–251.

Marcroft, S., Sprague, S., Pymer, S., Salisbury, P., and Howlett, B. (2004a). Crop isolation, not extended rotation length, reduces blackleg (leptosphaeria maculans) severity of canola (brassica napus) in south-eastern australia. Australian Journal of Experimental Agriculture, 44(6):601–606.

Marcroft, S., Sprague, S., Salisbury, P., and Howlett, B. (2004b). Potential for using host resistance to reduce production of pseudothecia and ascospores of leptosphaeria maculans, the blackleg pathogen of brassica napus. Plant Pathology, 53(4):468–474.

McCredden, J., Cowley, R. B., Marcroft, S. J., and Van de Wouw, A. P. (2018). Changes in farming practices impact on spore release patterns of the blackleg pathogen, Leptosphaeria maculans. Crop and Pasture Science, 69(1):1.

McGee, D. and Emmett, R. (1977). Black leg (leptosphaeria maculans (desm.) ces. et de not.) of rapeseed in victoria: crop losses and factors which affect disease severity. Australian Journal of Agricultural Research, 28(1):47–51.

Moghadam, P., Ward, D., Goan, E., Jayawardena, S., Sikka, P., and Hernandez, E. (2017). Plant disease detection using hyperspectral imaging. In Digital Image Computing: Techniques and Applications (DICTA), 2017 International Conference on, pages 1–8. IEEE.

Naseri, B., Davidson, J. A., and Scott, E. S. (2009). Maturation of pseudothecia and discharge of ascospores of Leptosphaeria maculans on oilseed rape stubble. European Journal of Plant Pathology, 125(4):523–531.

Otsu, N. (1975). A threshold selection method from gray-level histograms. Automatica, 11(285–296):23–27.

Pedregosa, F., Varoquaux, G., Gramfort, A., Michel, V., Thirion, B., Grisel, O., Blondel, M., Prettenhofer, P., Weiss, R., Dubourg, V., Vanderplas, J., Passos, A., Cournapeau, D., Brucher, M., Perrot, M., and Duchesnay, E. (2011). Scikit-learn: Machine learning in Python. Journal of Machine Learning Research, 12: 2825–2830.

Pound, M. P., Burgess, A. J., Wilson, M. H., Atkinson, J. A., Griffiths, M., Jackson, A. S., Bulat, A., Tzimiropoulos, G., Wells, D. M., Murchie, E. H., Pridmore, T. P., and French, A. P. (2016). Deep Machine Learning provides state-of-the-art performance in image-based plant phenotyping. Technical Report biorxiv;053033v1.

R Core Team (2018). R: A Language and Environment for Statistical Computing. R Foundation for Statistical Computing, Vienna, Austria.

Ren, S., He, K., Girshick, R., and Sun, J. (2017). Faster r-cnn: Towards real-time object detection with region proposal networks. IEEE Trans. Pattern Anal. Mach. Intell., 39(6):1137–1149.

Savage, D., Barbetti, M. J., MacLeod, W. J., Salam, M. U., and Renton, M. (2013). Temporal patterns of ascospore release in leptosphaeria maculans vary depending on geographic region and time of observation. Microbial ecology, 65(3):584–592.

Schindelin, J., Rueden, C. T., Hiner, M. C., and Eliceiri, K. W. (2015). The imagej ecosystem: an open platform for biomedical image analysis. Molecular reproduction and development, 82(7–8):518–529.

Simko, I., Jimenez-Berni, J. A., and Sirault, X. R. (2016). Phenomic approaches and tools for phytopathologists. Phytopathology, 107(1):6–17.

Sommer, C., Straehle, C., Kthe, U., and Hamprecht, F. A. (2011). Ilastik: Interactive learning and segmentation toolkit. In 2011 IEEE International Symposium on Biomedical Imaging: From Nano to Macro, pages 230–233.

Stewart, E. L., Hagerty, C. H., Mikaberidze, A., Mundt, C. C., Zhong, Z., and McDonald, B. A. (2016). An improved method for measuring quantitative resistance to the wheat pathogen zymoseptoria tritici using high-throughput automated image analysis. Phytopathology, 106(7):782–788.

Taylor, A., Coventry, E., Handy, C., West, J. S., Young, C. S., and Clarkson, J. P. (2018). Inoculum potential of sclerotinia sclerotiorum sclerotia depends on isolate and host plant. Plant Pathology.

The GIMP team (1997–2018). GNU image manipulation program. Available at www.gimp.org.

van der Walt, S., Schönberger, J. L., Nunez-Iglesias, J., Boulogne, F., Warner, J. D., Yager, N., Gouillart, E., Yu, T., and the scikit-image contributors (2014). scikit-image: image processing in Python. PeerJ, 2:e453.

Wang, G., Sun, Y., and Wang, J. (2017). Automatic image-based plant disease severity estimation using deep learning. Computational intelligence and neuroscience, 2017.

West, J. S., Kharbanda, P., Barbetti, M., and Fitt, B. D. (2001). Epidemiology and management of leptosphaeria maculans (phoma stem canker) on oilseed rape in australia, canada and europe. Plant Pathology, 50(1):10–27.

Wherrett, A., Sivasithamparam, K., and Barbetti, M. (2003). Chemical manipulation of leptosphaeria maculans (blackleg disease) pseudothecial development and timing of ascospore discharge from canola (brassica napus) residues. Australian Journal of Agricultural Research, 54(9):837–848.

Wherrett, A. D., Sivasithamparam, K., and Barbetti, M. J. (2004). Establishing the relationship of ascospore loads with blackleg (Leptosphaeria maculans) severity on canola (Brassica napus). Australian Journal of Agricultural Research, 55(8):849.

Yang, X. and Hong, C. (2018). Microsclerotial enumeration, size, and survival of calonectria pseudonaviculata. Plant Disease, 102(5):983–990.

Zhou, Z.-H. (2012). Ensemble Methods: Foundations and Algorithms. Chapman & Hall/CRC, 1st edition.

